# Multimerization- and glycosylation-dependent receptor binding of SARS-CoV-2 spike proteins

**DOI:** 10.1101/2020.09.04.282558

**Authors:** Kim M. Bouwman, Ilhan Tomris, Hannah L. Turner, Roosmarijn van der Woude, Gerlof P. Bosman, Barry Rockx, Sander Herfst, Bart L. Haagmans, Andrew B. Ward, Geert-Jan Boons, Robert P. de Vries

## Abstract

Receptor binding studies using recombinant SARS-CoV proteins have been hampered due to challenges in approaches creating spike protein or domains thereof, that recapitulate receptor binding properties of native viruses. We hypothesized that trimeric RBD proteins would be suitable candidates to study receptor binding properties of SARS-CoV-1 and -2. Here we created monomeric and trimeric fluorescent RBD proteins, derived from adherent HEK293T, as well as in GnTI mutant cells, to analyze the effect of complex vs high mannose glycosylation on receptor binding. The results demonstrate that trimeric fully glycosylated proteins are superior in receptor binding compared to monomeric and immaturely glycosylated variants. Although differences in binding to commonly used cell lines were minimal between the different RBD preparations, substantial differences were observed when respiratory tissues of experimental animals were stained. The RBD trimers demonstrated distinct ACE2 expression profiles in bronchiolar ducts and confirmed the higher binding affinity of SARS-CoV-2 over SARS-CoV-1. Our results show that fully glycosylated trimeric RBD proteins are attractive to analyze receptor binding and explore ACE2 expression profiles in tissues.

## INTRODUCTION

SARS-CoV-2 has sparked a society changing pandemic, and additional means to understand this virus will facilitate counter-measures. SARS coronaviruses carry a single protruding envelope protein, called spike, that is essential for binding to and subsequent infection of the host cell. The trimeric spike protein consists of 3 protomers of 140kD each, containing 60 N-linked glycosylation sites. The spike is composed of an S1 and S2 domain, in which S2 contains membrane fusion activity. S1 of coronaviruses can be further divided into N-terminal and C-terminal domains (NTD & CTD), both can contain the receptor-binding domain. In SARS-CoV-1 and -2 this domain is located in the CTD and referred to as the receptor-binding domain (RBD). The RBD binds to angiotensin-converting enzyme 2 (ACE2) [1-3], which functions as an entry receptor for SARS-CoV. After binding and internalization, several proteinases induce the spike protein into its fusogenic form allowing the fusion of the viral and target membrane. Although this pathway is known, the details that are of importance for receptor binding and what differentiates SARS-CoV-2 from its predecessor SARS-CoV-1, are incompletely understood. It has been shown that the affinity of the SARS-CoV-2 spike to ACE2 is significantly higher compared to SARS-CoV-1 [2, 4]. However, how this relates to tissue and cell tropism remains to be determined.

Several recombinant protein approaches to create coronavirus spike proteins have been utilized with success for the development of serological assays [5], elucidating of spike structures, and isolation of neutralizing antibodies [6, 7]. However, the conformation of the receptor-binding domain of the spike, which can be in the “up” and “down” configuration [8-12], is highly variable. This variability is important in receptor binding as only in the “up” conformation can RBD bind ACE2. Approaches to control the conformation for vaccine purposes using stabilization mutations appear to keep the RBD conformation in their down-state [11, 12], making them non preferred proteins to analyze receptor-binding properties. Also, the multimerization status of the RBD is critical to allow the analysis of ACE2 interactions. While monomeric RBD proteins can be efficiently made in large quantities [5], and bind ACE2, they are hardly used in receptor binding assays to cells and tissues. The majority of studies analyzing RBD protein binding to cells and tissues, utilize Fc-tagged proteins [2]. Fc-tags on recombinant proteins are extremely convenient for mammalian cell expression and purification systems, which result in dimeric recombinant spikes that are biologically functional. However, the coronavirus spike is trimeric and thus Fc tagged spikes do not fully recapitulate native properties.

We hypothesized that fluorescent trimeric RBD proteins would provide complementary means to study RBD-receptor interactions. In this study, we compared monomeric and trimeric SARS-CoV RBD with full-length trimeric spikes expressed in cells producing proteins with either complex or high mannose glycosylation. The fusion of sfGFP and mOrange2 at the C-terminus has previously been shown to increase expression yields, protein stability, and afford additional means for fluorescent-based experiments [13], and thus are attractive to be fused to RBD proteins. The resulting proteins were analyzed for binding to cell culture cells and paraffin-embedded tissues of various hosts including susceptible and non-susceptible animals. The results demonstrate that fully glycosylated trimeric SARS-CoV-2 RBD proteins reveal the differences in ACE2 expression between cell cultures and tissue sections. These trimeric RBD proteins bind ACE2 efficiently in a species-dependent manner and can be used to profile ACE2 tissue expression. Finally, we observed distinct expression of ACE2 in bronchioles of experimental animal models.

## RESULTS

### Generation of fluorescent coronavirus receptor-binding domain spike proteins

To create recombinant fluorescent soluble full-length ectodomains, NTDs and RBDs, we cloned these open reading frames (ORF) in plasmids with and without the GCN4 trimerization domain, fused to either sfGFP or mOrange (Fig 1A). The monomeric and trimeric RBDs were efficiently expressed in both HEK 293T as well as GnTI cells (data not shown), with an increased yield up to 2- to 5-fold when fused to a C-terminal sfGFP (Fig 1B). Expression yields of the mOrange2 fusions were comparable to that of sfGFP fusions (data not shown). To illustrate the expression yields of SARS spike proteins or domains thereof we measured the fluorescence in the cell culture supernatant (Fig 1C). The wild-type full-length ectodomains were difficult to express even with the addition of sfGFP or mOrange2 fusion (Fig 1B). To increase yields for the full-length ectodomain we introduced the 2P and additional hexapro mutations [14], and analyzed the fluorescence in cell culture supernatants five days post-transfection after incubation at 33 or 37°C. Although we did not observe a large increase in yields, we were able to purify sufficient protein to compare full-length ectodomain trimers vs monomeric and trimeric RBD and NTD proteins.

**Figure 1.**
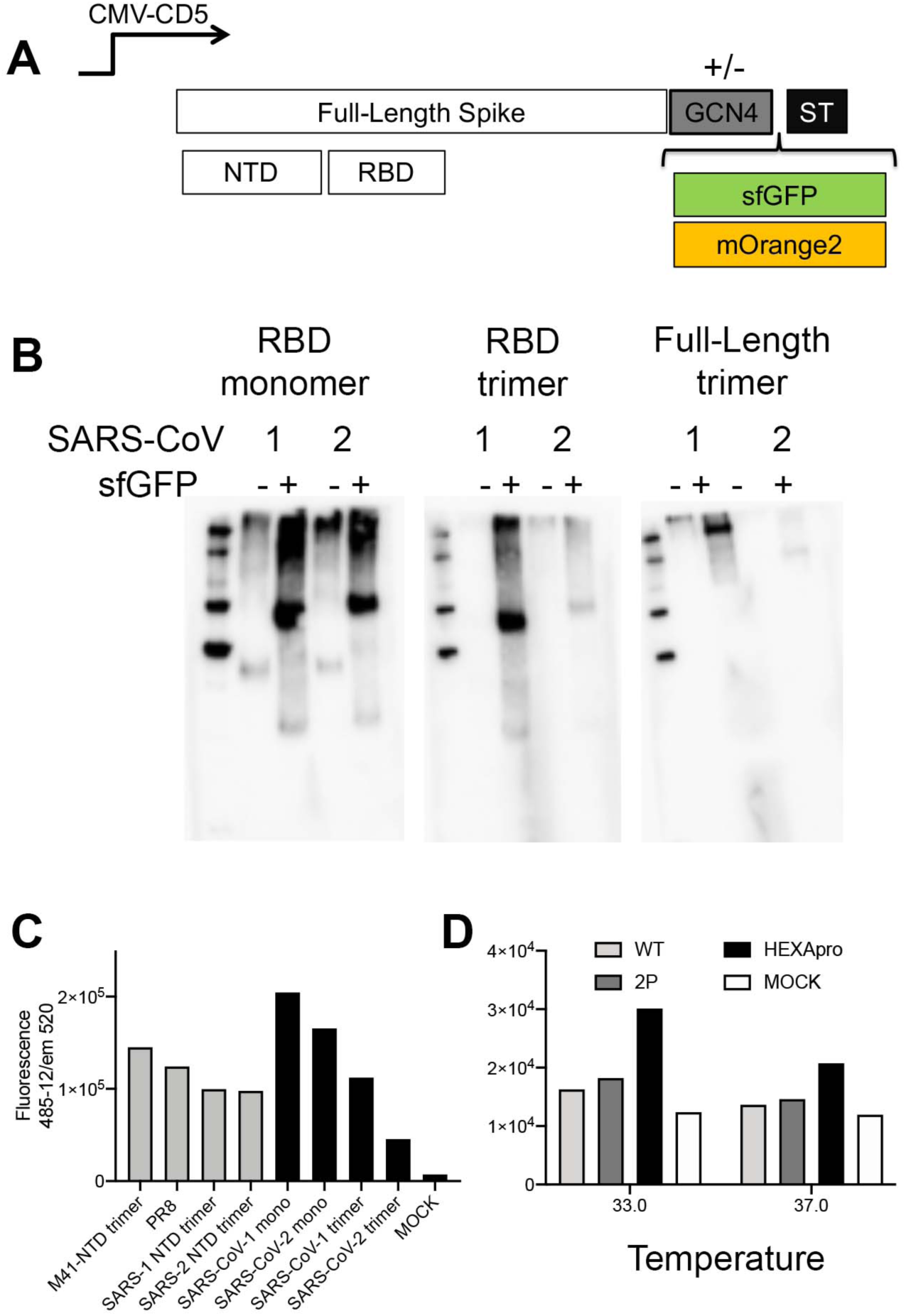
Expression of coronavirus spike proteins using sfGFP and mOrange2 fusions. **(A) Spike expression plasmids with and without GCN4 trimerization motif fused with either sfGFP or mOrange2** Schematic representation of the used spike expression cassette. The spike open reading frame is under the control of a CMV-promotor and was cloned in frame with DNA sequences coding for the CD5 signal peptide. At the C-terminus a GCN4 trimerization domain followed by sfGFP or mOrange2, and a TEV cleavable Strep-tag II. **(B) Expression analyses of non and sfGFP fused monomeric and trimeric RBD proteins and the full-length spike** Denatured samples of the cell culture supernatant were subjected to SDS-PAGE and western blot analyzes stained with an anti-Streptag-HRP antibody. **(C) Quantification** sfGFP emission was directly measured in the supernatants. **(D) Full-length spike adaptation to the hexapro variant**. Full-length spike expression vectors containing the wildtype, 2P or Hexapro were expressed at 33 or 37C and fluorescence was measured in cell culture supernatant 4 days post transfection.

### Spike RBD domains in frame with a C-terminal GCN4 and fluorescent reporter protein display multimeric features on gel and maintain antigenicity

After purification, all RBD proteins were analyzed on gel under non- and reducing conditions (Fig 2A). Without reducing agent, monomeric RBD proteins revealed dimeric fractions which could be reduced to a single monomeric form. The NTD trimers were reduced under non-reducing conditions, thus solely by SDS. The trimeric RBD variants, on the other hand, revealed dimers and trimers that could be reduced. Besides, the NTD of prototypical γ-coronavirus IBV-M41 and influenza A virus HA PR8 as control proteins were included. Finally, we determined the extent of N-glycosylation maturation on purified proteins expressed in either GnTI or 293T by subjecting the monomeric and trimeric proteins to PNGaseF and EndoH treatment (Fig S1). Upon PNGaseF treatment, all N-glycans were trimmed whereas 293T derived proteins were insensitive to Endo H, as expected. Surprisingly trimeric RBDs derived from GnTI cells were partially resistant to Endo H treatment, whereas the monomers derived from GnTI cells where fully deglycosylated using Endo H.

**Figure 2.**
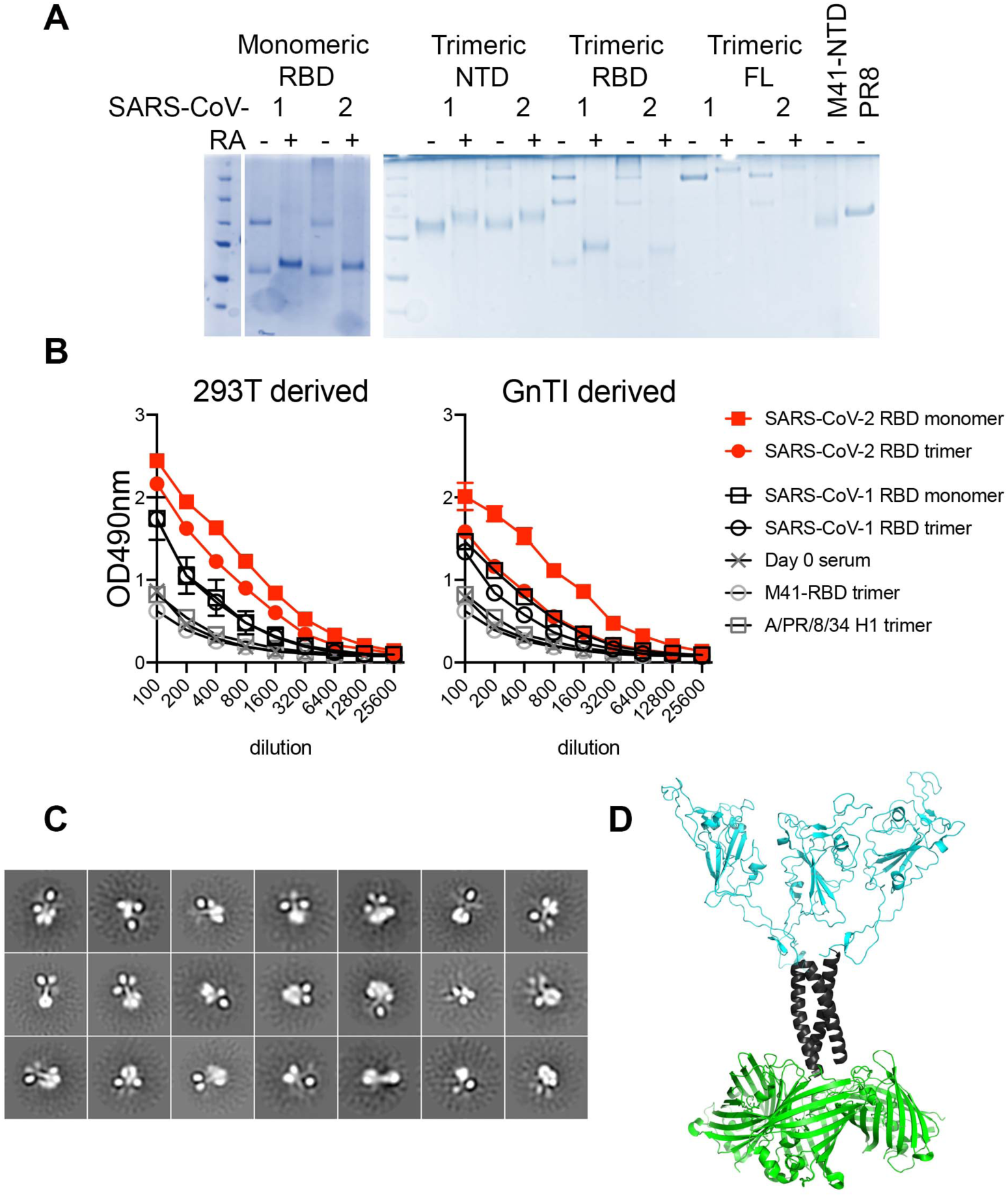
Molecular analyses. **(A) SDS-PAGE analyses of purified RBD proteins** 500 ng of purified proteins were loaded on a gel with or without preheating for 30 minutes at 98C in the presence of a reducing agent. **(B) Antigenic analyses using 21-day post-infection macaque serum** 2μg/ml SARS-CoV-RBD proteins were coated on 96-well plates. Proteins were detected with macaque serum for 2hrs at RT. An anti-human IgG-HRP was used to detect RBD specific antibodies. Both 293T and GnTI derived proteins were analyzed. As negative controls, IBV-M41-NTD and HA-PR8D were included. **(C) Negative stain EM of trimeric RBD fused to sfGFP** Negative-stain 2D class averages of soluble RBD proteins demonstrate that they are well-folded trimers. The C-terminal helices and fusion proteins are visible in some class averages. **(D) Structural model** Based on the crystal structure of a SARS-CoV-2 RBD in the up conformation, with a GNC4 trimerization domain with 3 sfGFP domains added.

### Fluorescent multimeric Spike RBD proteins maintain antigenicity

Next, we examined the antigenicity of the SARS-CoV-1 and -2 proteins using serum collected from macaques 21 days post-infection with SARS-CoV-2 [15]. Both SARS-CoV-2 RBD monomers and trimers derived from 293T cells were efficiently recognized, indicating proper folding (Fig 2B). As expected, SARS-CoV-1 RBD proteins were poorly recognized, and the negative controls M41 NTD and PR8 HA displayed baseline binding identical to pre-infection serum. The NTD trimers were likewise not recognized by the serum (not shown), indicating that the majority of antibodies in naïve animals after infection are directed against the SARS-CoV RBD [16]. Similar results were obtained using GnTI-derived proteins, with the RBD trimer being less efficiently recognized by the macaque serum than its monomeric counterpart. This is in line with recent observations that insect cell-derived proteins are less well bound by serum antibodies [5], indicating the importance of mature N-glycans.

### RBD fused to GCN4 and sfGFP fold as trimers with three RBD and sfGFP molecules divided by the trimerization coiled-coil

To determine whether the fluorescent RBD trimers are indeed structured in a trimeric manner we subjected these proteins to negative stain single-particle EM. The EM data revealed that the RBD proteins form stable trimers that resemble known spike structures (Fig. 2C). Initially, 58,018 individual particles were picked, placed into a stack, and submitted to reference-free two-dimensional (2C) classification. From the initial 2D classes, particles that did not resemble RBD were removed, resulting are final particle stacks of 32,152 particles, which were then subject to Relion 2D classification. All resultant classes demonstrated evident and distinct trimeric RBD, GCN4, and three sfGFP protein structures that could be identified in the EM images. From the EM images, we generated a model in which we took the crystal structures of sfGFP, the GCN4 trimerization domain (PDB:2O7H), and the SARS-CoV-2 RBD (PDB: 6XM4) to demonstrate the likely structure of our RBD trimer (Fig 2D).

### Fluorescent multimeric Spike RBD proteins bind cell lines in an ACE2 dependent manner and similar to the full-length ectodomain

To determine the biological activity of our RBD proteins we stained A549 and VERO cells that are reported to support SARS-CoV replication, with the latter being more susceptible [17]. However, A549 cells were bound by all our RBD proteins with a slight increase in intensity from monomeric GnTI derived RBD proteins to trimeric 293T derived RBDs (Fig 3). Trimeric 293T RBD binding was efficiently blocked using 4μM recombinant ACE2 whereas 400nM ACE2 pre-incubation was not sufficient to prevent binding of fully glycosylated trimeric RBD proteins to cells completely. SARS-CoV-2 RBD proteins bound slightly more intensely to A549 cells compared to the same SARS-CoV-1 RBD proteins. A similar pattern was observed for VERO-E6 cells, however, the fully glycosylated SARS-CoV-2 RBD trimer bound markedly stronger compared to the other RBD preparations (Fig S2A). Importantly, the full-length ectodomain also bound efficiently to A549 cells (Fig S2B). We did not observe any binding of the trimeric NTD domains to A549 cells (Fig S2B). MDCK cells, derived from canine kidney, served as negative controls, to which we indeed did not observe any binding with any of the indicated proteins (Fig S2C).

**Figure 3.**
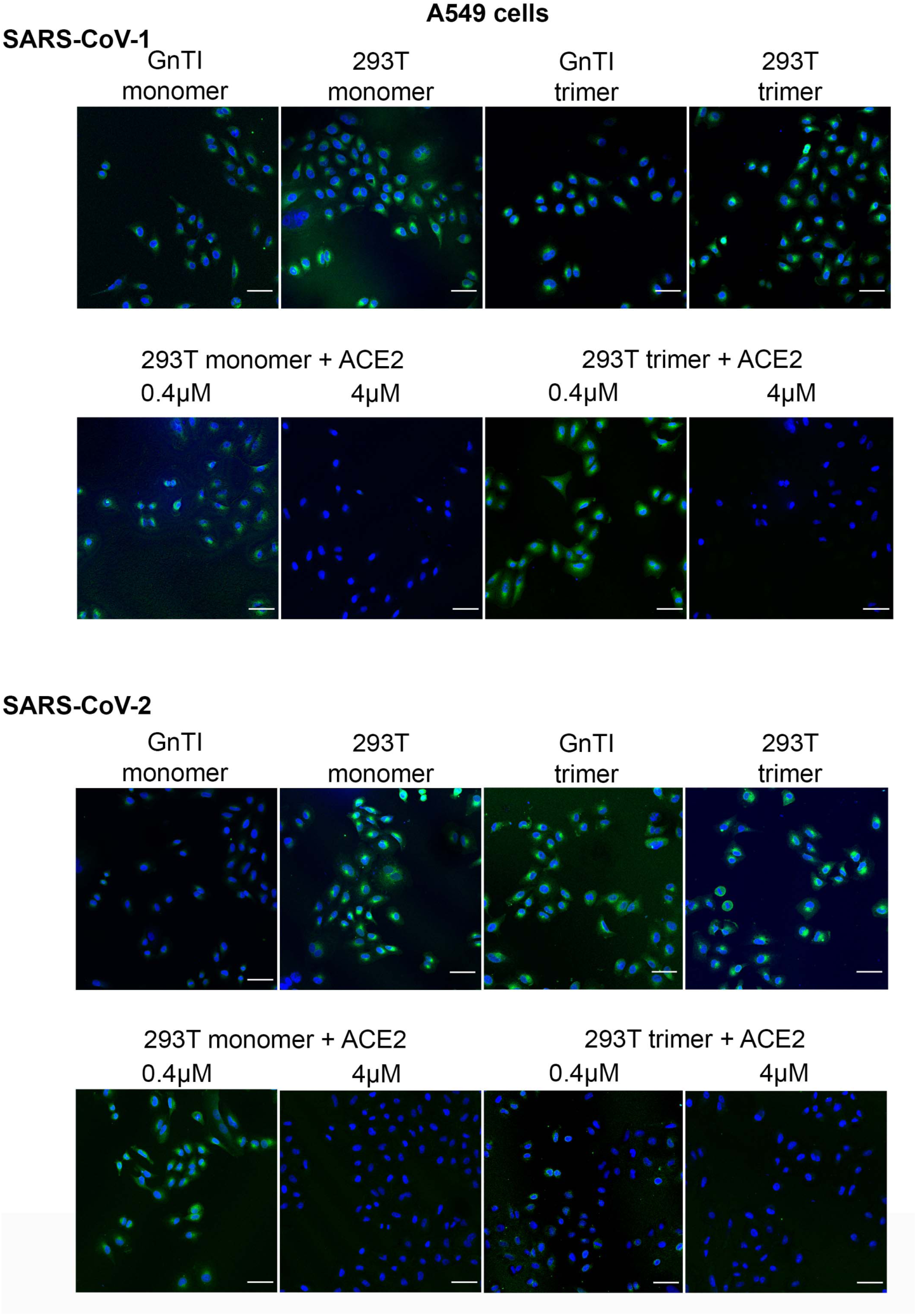
Binding of RBD proteins to A549 cells. SARS-CoV-RBD proteins were applied at 50μg/ml onto A549 cells and where indicated pre-incubated with recombinant ACE2 protein. SARS-CoV proteins were detected using an anti-Streptag and a goat-anti-mouse antibody sequentially. sfGFP fused SARS-CoV-RBD proteins were applied from GnTI monomers to 293T trimers, left to right. Scalebar is 50μm.

### Fluorescent RBD proteins reveal strong binding to bronchioles in both ferret and Syrian golden hamster lungs

Ferrets are a susceptible animal model for SARS-CoV-2 [18, 19] and closely related minks are easily infected on farms [20]. Formalin-fixed, paraffin-embedded tissue slides, will most closely resemble the complex membrane structures to which spike proteins need to bind. First, ACE2 expression was assessed using an ACE2 antibody which allowed for comparisons with SARS-CoV-RBD protein binding localization. Here, ACE2 expression was observed in bronchiolar epithelium cells of ferret lungs (Fig 4A), while the antibody failed to bind mouse lungs. Comparably, the SARS-CoV RBD protein preparations did not stain mouse lungs. In ferret lung tissues, we observed only minimal binding of monomeric RBD proteins derived from GnTI cells (Fig 4B). The fully glycosylated counterparts showed a slight increase in fluorescent intensity, which was further increased when fully glycosylated trimeric RBDs were applied. In all cases, SARS-CoV-2 displayed a higher avidity compared to SARS-CoV-1. A similar trend of binding intensities was observed for monomeric, trimeric, and different N-glycosylated SARS-CoV-RBD proteins fused to mOrange2 (Fig S3A). Again specific binding was seen to the epithelium of terminal bronchioles and, to a much lower extent, to alveoli and endothelium. The results were confirmed using horseradish peroxidase readout with a hematoxylin counterstain (Fig S3B), which output is enzyme driven and purely qualitative, however, we did observe similar differences in staining intensities. Here, very minimal staining using the SARS-CoV NTD domains was observed (Fig S3B), which we did not detect using a fluorescent readout (Fig S3C). To determine if the binding was ACE2 dependent we pre-incubated trimeric RBD proteins with recombinant ACE2. While 4 μM was sufficient to block binding to cell culture cells (Fig 4C), 8μM was needed to prevent all detectable binding to ferret lung tissue.

**Figure 4.**
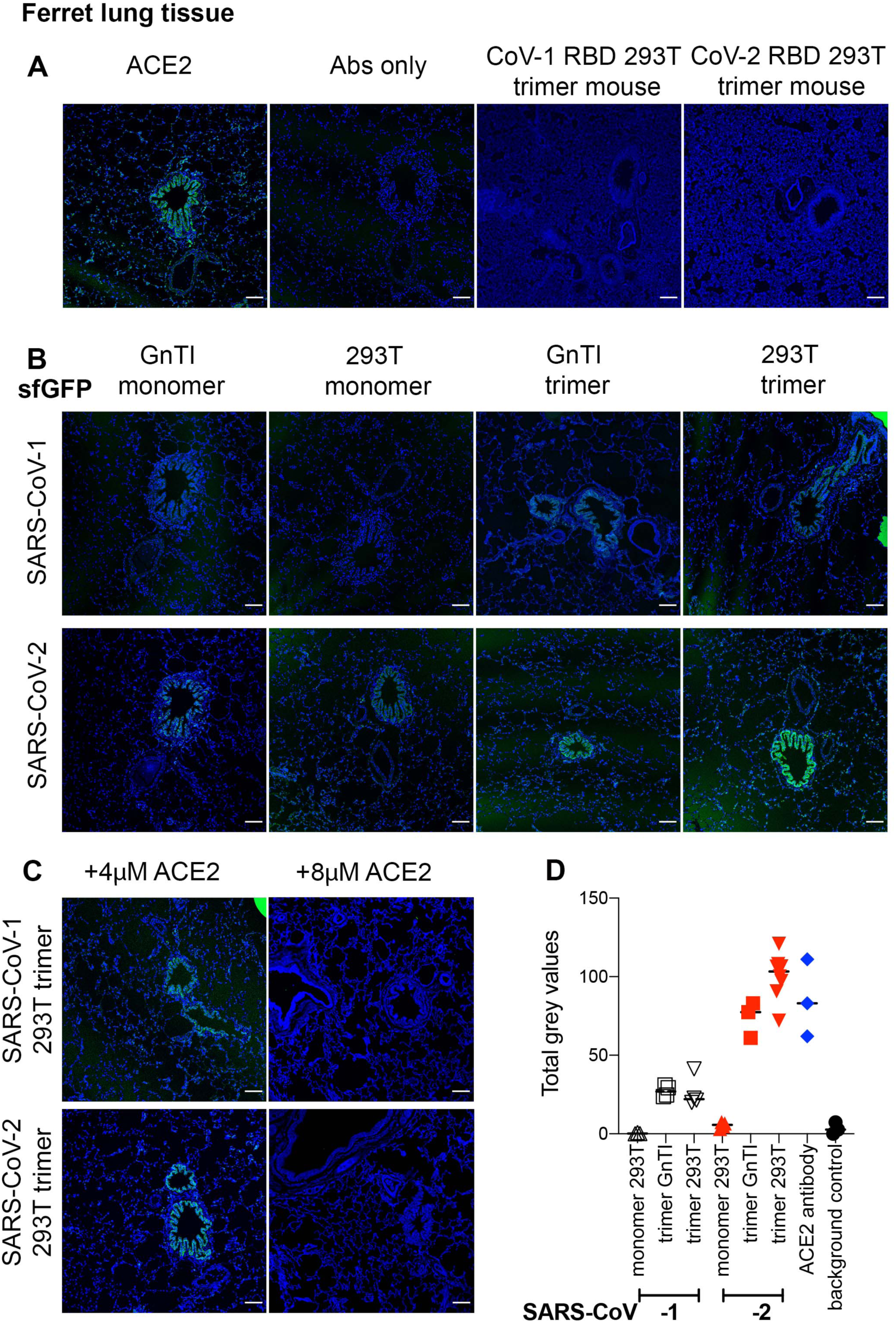
Binding of SARS-COV-RBD proteins and ACE2 antibody to mouse and ferret lung tissue slides. **(A) ACE2 antibody, antibody control, and SARS-CoV-1 and -2 on mouse lung**. Scalebar is 100μm. **(B) SARS-CoV-RBD fluorescent protein localization in ferret lung tissue**. SARS-CoV-RBD proteins were applied at 50μg/ml and detected using an anti-Streptag and goat-anti-mouse antibodies sequentially. DAPI was used as a nucleic stain. Scalebar is 100μm. **(C) SARS-CoV-RBD trimer produced in HEK 293T cells pre-incubated with recombinant ACE2 before application on ferret lung tissue slides**. **(D) Quantification** The intensity of gray pixels of stained ferret lung tissue slides with ImageJ version 1.52p.

**Figure 5.**
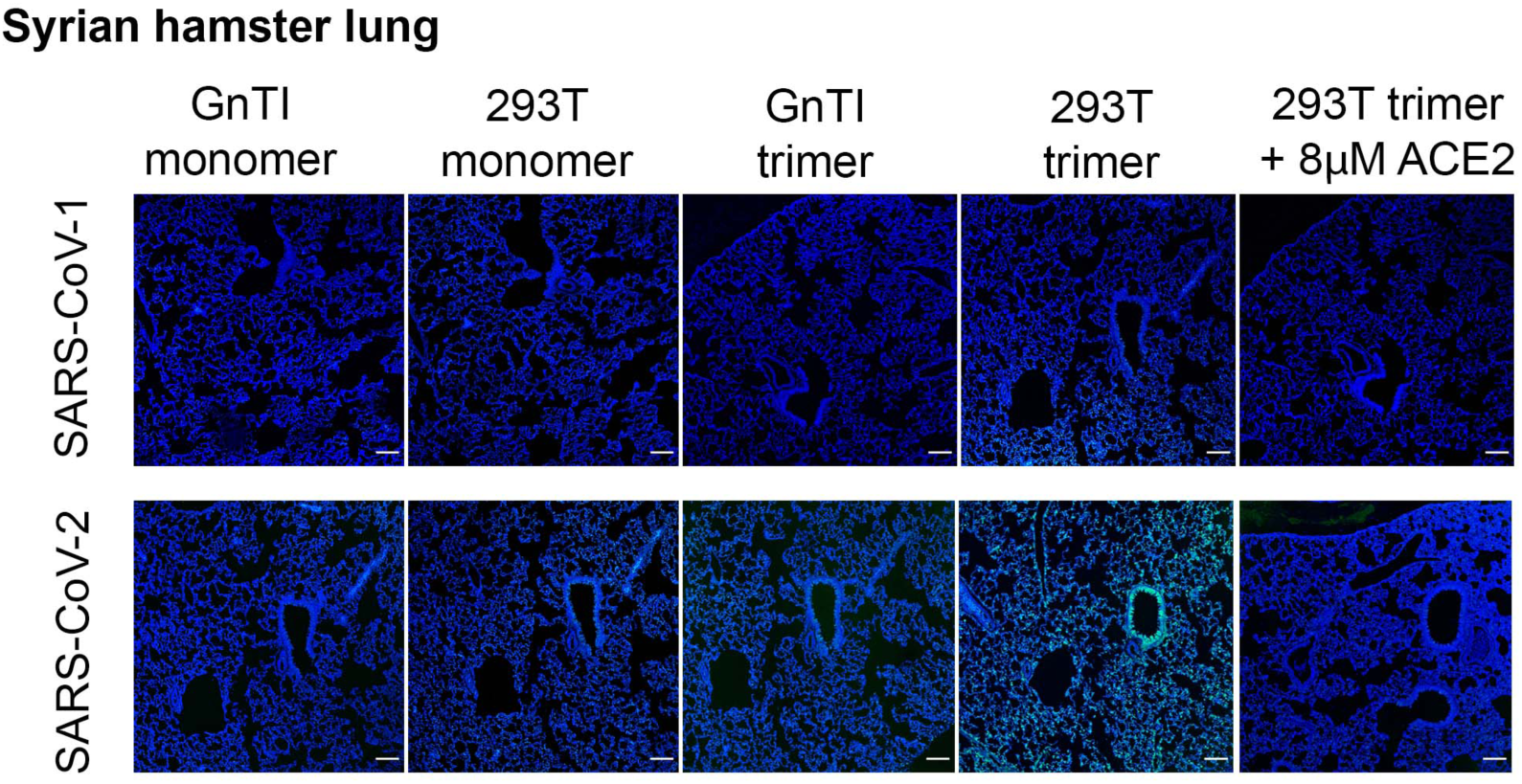
Binding of SARS-CoV-RBD proteins to Syrian hamster tissue slides. SARS-COV-RBD proteins (50μg/ml) were applied to Syrian hamster lung tissue slides and detected using additional anti-Streptag and goat-anti-mouse antibodies sequentially. Where indicated SARS-CoV-RBD proteins were pre-incubated with recombinant ACE2 protein. Scalebar is 100μM.

To confirm our observations of different binding on tissues, we quantified the intensities of the ACE2 antibody and SARS-CoV-1 and -2 RBD proteins, except for the monomeric GnTI derived proteins as these were almost at the background (Fig 4D). As expected a noteworthy trend was observed of increasing binding strength from SARS-CoV1 GnTI derived monomers to SARS-CoV-2 fully glycosylated RBD trimers. Interestingly multimerization appears to be more important for strong ACE2 interaction to tissue compared to the glycosylation status.

Finally, we applied our RBD proteins on the lung tissue of Syrian hamsters, which are currently the preferred small animal model and known to be highly susceptible to SARS-CoV-1 and -2 [21-23]. Here again, trimeric fully glycosylated RBD proteins were superior in binding compared to their monomeric counterparts. Interestingly, SARS-CoV-2 bound with an even higher intensity to terminal bronchioles compared to SARS-CoV-1. Importantly, all binding could be inhibited by pre-incubation of the RBD proteins with recombinant ACE2, indicating that binding intensity differences are likely ACE2 related.

## DISCUSSION

In this report, we describe the generation of fluorescent trimeric RBD proteins derived from SARS-CoV-1 and 2. These proteins can be efficiently expressed in adherent cells and purified directly from the cell culture supernatant using the increased protein yields provided by the introduction of a C-terminal sfGFP or mOrange fusion. The fluorescent RBD trimers were superior in receptor binding on tissues compared to their monomeric RBD counterparts.

Mature N-glycosylation on the spike protein appears to be important for the avid binding of trimeric RBD proteins to ACE2 in different assays. There is a wealth of information on site-specific N- and O-glycan conformations on spike proteins [24-26], with some disagreement on the nature of the RBD N-glycans. In our protein preparations, they appear to be complex as the majority could not be cleaved by EndoH (Fig S1). Furthermore, the N-glycosylation of the NTD domain has been shown to affect the conformation of the RBD in full-length spikes [27], however similar analyses on the RBD N-glycans are lacking.

Currently, most knowledge of ACE2 expression is based on genomic and transcriptomic data [28]. However, these analyses are limited as it does not determine expression biochemically on epithelial cells that make contact to the outside world [29]. Several groups have transfected variants of ACE2 in cells to analyze transduction by SARS-CoV-pseudo viruses, although informative these studies do not provide information on the natural expression pattern of ACE2 in susceptible hosts [30, 31]. A detailed understanding of the difference between animal species and cell-specific expression of ACE2 at the molecular level is essential, as this can provide valuable knowledge on potential hosts that can be susceptible to SARS-like coronaviruses. Here we demonstrate that our fluorescent trimeric RBD proteins are complementary to study ACE2 expression and SARS-CoV receptor binding dynamics.

An intriguing observation was the abundant expression of ACE2 in terminal bronchioles both in the ferret and Syrian hamster lungs, which are used as experimental animal models. SARS-CoV-RBD protein binding correlated perfectly with the staining of the ACE2 antibody and a ferret infection study [18]. In another ferret infection study, however, the infection was observed in nasal epithelium and many type I pneumocytes and fewer type II pneumocytes in the lungs, with few bronchial epithelial cells expressing viral antigen [19]. Some studies have been performed with anti-ACE2 antibodies, but these stainings appear not to correlate with infection patterns [32, 33]. We observed a similar trend indicating that several other factors may be important to infect cells and that infection could, in turn, regulate ACE2 expression.

## MATERIAL AND METHODS

### Expression and purification of coronavirus spike proteins for binding studies

Recombinant SARS-CoV-1 and -2 envelope proteins and their subunits were cloned using Gibson assembly from cDNAs encoding codon-optimized open reading frames of full-length SARS-CoV-1 and -2 spikes (A kind gift of Rogier Sanders, Amsterdam Medical Centre, The Netherlands). The pCD5 expression vector as described previously [34], was adapted to clone the SARS-1 (GenBank: MN908947.3) and 2 (GenBank: MN908947.3), ectodomains (SARS-2 15-1213, SARS-1 15-1195), N-terminal S1 (SARS-2 15-318, SARS-1 15-305, M41 19 to 272) and RBDs (SARS-2 319-541, SARS-1 306-527) sequences coding for a secretion signal sequence, a GCN4 trimerization domain (RMKQIEDKIEEIESKQKKIENEIARIKK) followed by a seven amino acid cleavage recognition sequence (ENLYFQG) of tobacco etch virus (TEV), a super folder GFP [13], or mOrange2 [35] and the Twin-Strep (WSHPQFEKGGGSGGGSWSHPQFEK); IBA, Germany). Alongside we expressed infectious bronchitis virus M41 spike RBD (GenBank: AY851295.1) [36],) and influenza A virus PR8 HA (GenBank: NP_040980) [13]. The viral envelope proteins were purified from cell culture supernatants after expression in HEK293T or HEK293GnTI(-) cells as described previously [34]. In short, transfection was performed using the pCD5 expression vectors and polyethyleneimine I. The transfection mixtures were replaced at 6 h post-transfection by 293 SFM II expression medium (Gibco), supplemented with sodium bicarbonate (3.7 g/L), Primatone RL-UF (3.0 g/L), glucose (2.0 g/L), glutaMAX (Gibco), valproic acid (0,4 g/L) and DMSO (1,5%). At 5 to 6 days after transfection, tissue culture supernatants were collected.

For ACE2 inhibition studies, ACE2 was expressed in a highly identical fashion, ACE2 (Addgene Plasmid #145171) was cloned into a pCD5 expression vector with SacI and BamHI restriction enzymes. The adapted pCD5 expression vector with an N-terminal HA leader (MKTIIALSYIFCLVFA) peptide, ACE2, and Twin-Strep (WSHPQFEKGGGSGGGSWSHPQFEK); IBA, Germany) was purified from HEK293T cell culture supernatant.

### Determining expression yield

We measure fluorescence in the cell culture supernatant when applicable using a polarstar fluorescent reader with excitation and emission wavelengths of 480 nm and 520 nm for sfGFP and 520nm and 550nm for mOrange2, respectively. Spike protein expression was confirmed by western blotting using a StrepMAB-HRP classic antibody. Proteins are purified using a single-step with strepTactin sepharose beads in batch format. Purified proteins were pre-treated (were indicated) with PNGase F or Endo H (New England Biolabs, USA) according to the manufacturer’s protocol before analysis by Western blotting using Strep-MAb-HRP classic antibody.

### Antigen ELISA

Plates were coated with 2 μg/mL spike protein in PBS for 16 hours at 4°C, followed by blocking by 3% bovine serum albumin (BSA, VWR, 421501J) in phosphate-buffered saline-Tween 0,1% (PBS-T 0,1%). After the block, the proteins were detected using a 21-days post-infection macaque serum and serum of the same monkey before infection. Serum starting dilution was 1:100 and diluted 1:1 for 5 times and incubated at RT for 2 hrs. Next, wells were treated with goat-α-human HRP secondary antibody for 1 hour at room temperature. Serum spike binding antibodies were detected using ODP and measured in a plate reader (Polarstar Omega, BMG Labtech) at 490 nm.

### Negative stain electron microscopy structural analysis

SARS-CoV2-RBD-GCN4-sfGFP in 10mM Tris, 150mM NaCl at 4°C was deposited on 400 mesh copper negative stain grids and stained with 2% uranyl formate. The grid was imaged on a 200KeV Tecnai F20 electron microscope and a 4k x 4k TemCam F416 camera. Micrographs were collected using Leginon [37] and then uploaded to Appion [38]. Particles were picked using DoGPicker [39], stacked. 2D processing was undertaken using Relion. Images showing Trimeric RBD, a GCN4 connector, and three sfGFPs.

### Immunofluorescent cell staining

Vero-E6, A549, and MDCK cells grown on coverslips were analyzed by immunofluorescent staining. Cells were fixed with 4% paraformaldehyde in PBS for 25 min at RT after which permeabilization was performed using 0,1% Triton in PBS. Subsequently, the coronavirus spike proteins were applied at 50μg/ml for 1 h at RT. Primary Strep-MAb classic chromeo-488 (IBA) and secondary Alexa-fluor 488 goat anti-mouse (Invitrogen) were applied sequentially with PBS washes in between. DAPI (Invitrogen) was used as nuclear staining. Samples were imaged on a Leica DMi8 confocal microscope equipped with a 10x HC PL Apo CS2 objective (NA. 0.40). Excitation was achieved with a Diode 405 or white light for excitation of Alexa488 and Alexa555, a pulsed white laser (80MHz) was used at 488 nm and 549 nm, and emissions were obtained in the range of 498-531nm and 594-627 nm respectively. Laser powers were 10 – 20% with a gain of a maximum of 200. LAS Application Suite X was used as well as ImageJ for the addition of the scale bars.

### Tissue staining

Sections of formalin-fixed, paraffin-embedded ferret, Syrian hamster, and mouse lungs were obtained from the Department of Veterinary Pathobiology, Faculty of Veterinary Medicine, Utrecht University and the department of Viroscience, Erasmus University, The Netherlands, respectively. Tissue sections were rehydrated in a series of alcohol from 100%, 96% to 70%, and lastly in distilled water. Tissues slides were boiled in citrate buffer pH 6.0 for 10 min at 900 kW in a microwave for antigen retrieval and washed in PBS-T three times. Endogenous peroxidase activity was blocked with 1% hydrogen peroxide for 30 min. Tissues were subsequently incubated with 3% BSA in PBS-T overnight at 4 °C. The next day, the purified viral spike proteins (50μg/ml) were added to the tissues for 1 h at RT. With rigorous washing steps in between the proteins were detected with mouse anti-strep-tag-antibodies (IBA) and goat anti-mouse IgG antibodies (Life Biosciences). Where indicated recombinant RBD and ACE2 proteins were pre-incubated O/N at 4 °C before application on lung tissues or cells.

### Quantifying and plotting fluorescent image data

Image quantification was performed by measuring the intensity of gray pixels of the stained ferret and Syrian hamster lung tissue slides with ImageJ version 1.52p. For stained tissues, background correction was performed by subtracting the average signal intensity of the antibody control from the images stained with ACE2 antibody, SARS-CoV1, and SARS-CoV2 recombinant proteins. Regions of interest were set by highlighting the area using the Image -> Adjust -> Threshold setting. The 8-bit images were checked for saturation by plotting the distribution of gray values with Analyze -> Histogram. Subsequent analysis was performed by quantifying the mean ± standard deviation of alveoli/cells stained with distinct proteins in various images (Analyze -> Measure). Plotting means ± standard deviation values were performed in GraphPad Prism v8.0.1. A 3D Surface plot was generated for several staining conditions with Analyze -> 3D Surface Plot.

## ACKNOWLEDGEMENTS

R.P.dV is a recipient of an ERC Starting Grant from the European Commission (802780) and a Beijerinck Premium of the Royal Dutch Academy of Sciences. Electron microscopic imaging work in the Ward lab was supported by Bill and Melinda Gates Foundation OPP1170236. SH was funded by NIH/NIAID (contract number HHSN272201400008C). The authors would like to thank Rogier Sanders and Phillip Brouwer of the Amsterdam Medical Centre for sharing the SARS-CoV 1 & 2 open reading frames, Andrea Gröne, Darryl Leydekkers, and Hélène Verheije from the Division of Pathology, Department Biomolecular Health Sciences, Faculty of Veterinary Medicine, Utrecht University, for sharing tissues, IBV M41 plasmids and editing of the manuscript.

---

**Supplemental figure 1.**
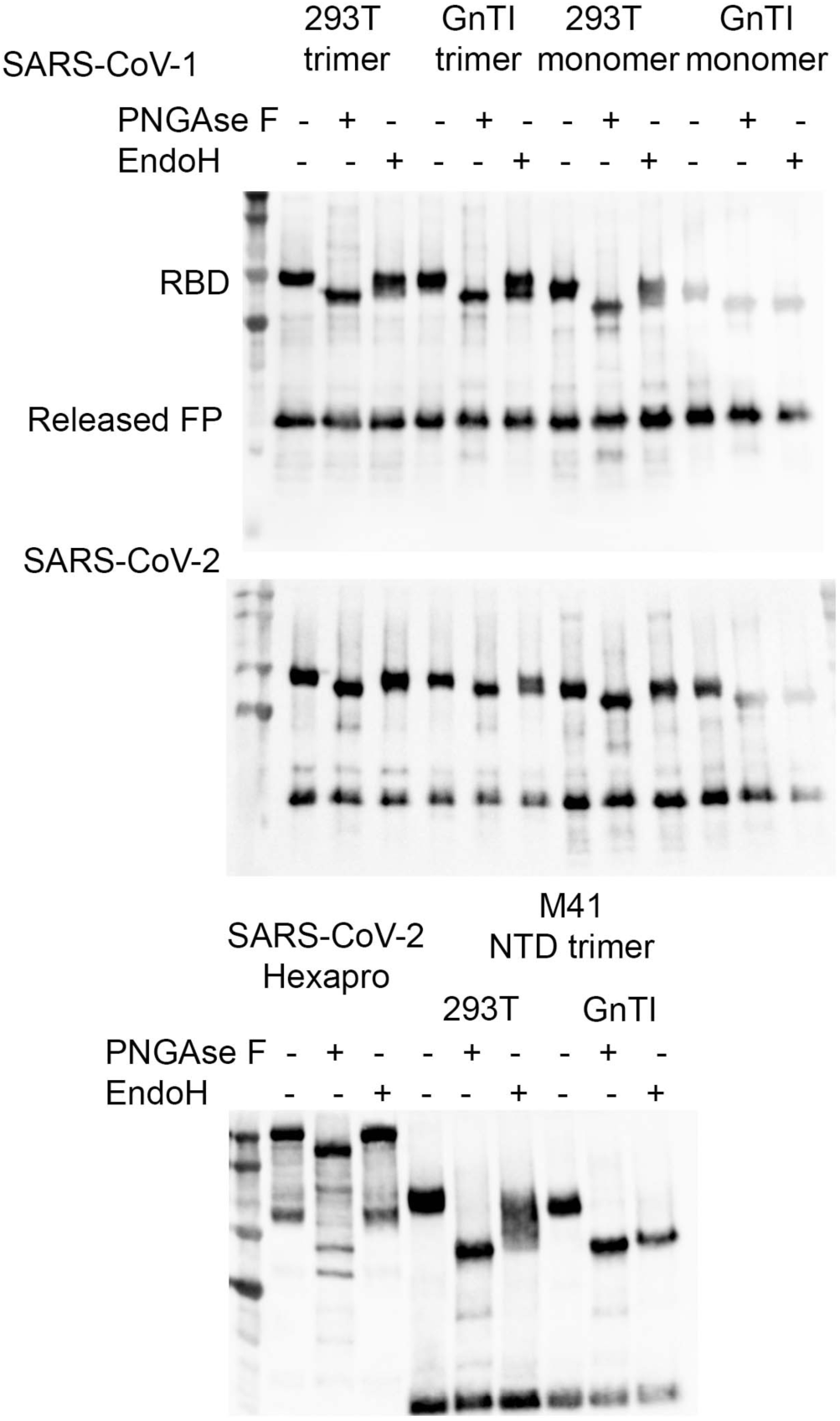
Glycosylation analyses of the SARS-CoV protein preparations. 0.5μg of protein was subjected without or with PNGase F or EndoH for 1hr and subjected to SDS-PAGE and western blot analyzes.

**Supplemental figure 2.**
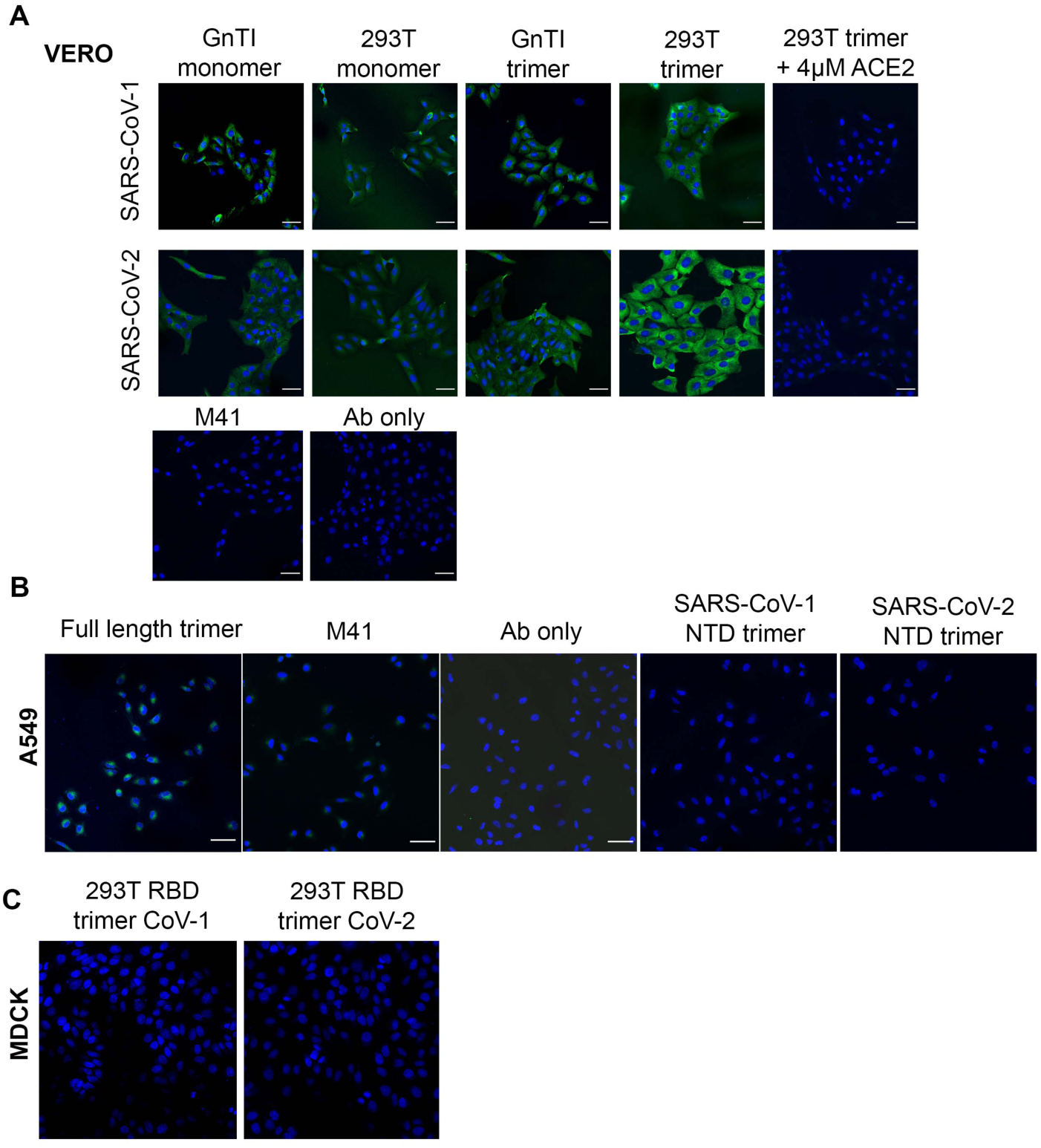
Binding of RBD proteins to cell lines. **(A) Protein binding of RBD proteins observed on VERO E6 cells**. Proteins were applied 50μg/ml and where indicated pre-incubated with recombinant ACE2 protein. Spike proteins were detected using anti-strep and goat-anti-mouse antibodies. Scalebar is 5μm. **(B) Binding of full-length SARS-CoV-2 ectodomain, IBV-M41, antibodies only, and NTD spike proteins to A549 cells**. Proteins were applied 50μg/ml and detected using anti-strep and goat-anti-mouse antibodies. Scalebar is 50μm. **(C) Non-binding of RBD trimers to MDCK cells**.

**Supplemental figure 3.**
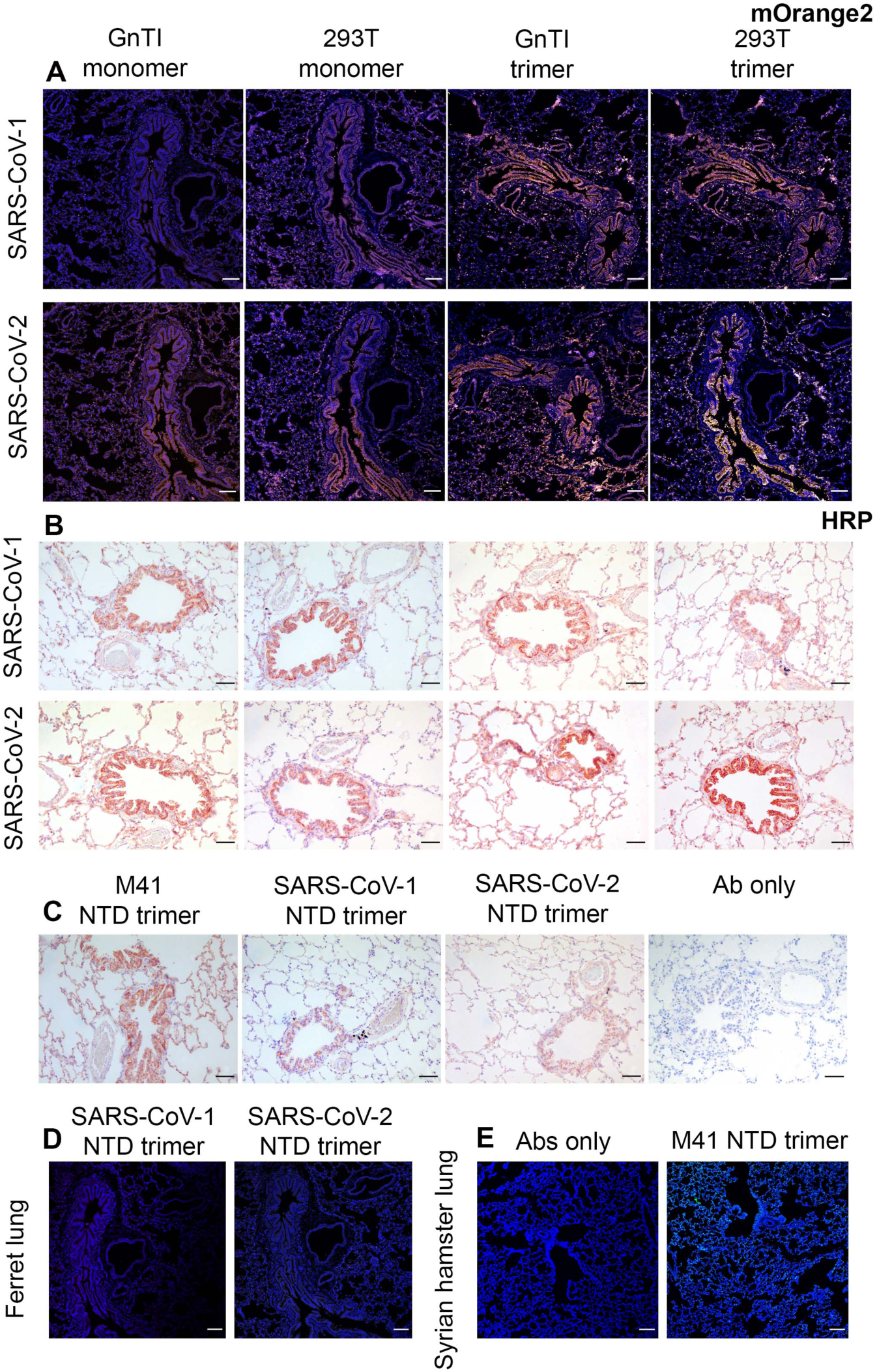
Binding of RBD proteins to tissues. **(A) Binding of RBD proteins fused to mOrange2 to ferret lung tissues** Proteins were applied 50μg/ml and detected using anti-strep and goat-anti-mouse antibodies. Scalebar is 100μm. **(B) Binding of RBD proteins fused to sfGFP proteins to ferret lung tissues, using HRP as a readout**. Identical experiment to (A) but using an HRP readout using anti-strep and goat-anti-mouse antibodies. Scalebar is 100μm. **(C) Control staining on ferret lung tissues using HRP as readout**. M41, NTD trimers of SARS-CoV-1 and -2 and antibodies only. Proteins were applied 50μg/ml and detected using anti-strep and goat-anti-mouse antibodies. Scalebar is 100μm. **(D) Lack of NTD binding to ferret lung tissue using fluorescence** Proteins were applied 50μg/ml and detected using anti-strep and goat-anti-mouse antibodies. Scalebar is 100μm. **(E) Control stainings to Syrian hamster tissues, antibodies only and M41** Proteins were applied 50μg/ml and where indicated pre-incubated with recombinant ACE2 protein. Scalebar is 100μm.

